# Replication Challenges in Linking Personality to Resting-State Functional Connectomics

**DOI:** 10.64898/2026.01.19.700331

**Authors:** Nikola Jajcay, David Tomeček, Iveta Fajnerová, Jan Rydlo, Jaroslav Tintěra, Jiří Horáček, Jiří Lukavský, Jaroslav Hlinka

**Affiliations:** Department of Complex Systems, Institute of Computer Science, Czech Academy of Sciences, Prague, Czech Republic; National Institute of Mental Health, Klecany, Czech Republic; Department of Radiology, Institute for Clinical and Experimental Medicine (IKEM), Prague, Czech Republic; Institute of Psychology, Czech Academy of Sciences, Prague, Czech Republic

**Keywords:** personality neuroscience, functional connectivity, graph theory, Big Five, fMRI

## Abstract

An increasing number of studies are currently focusing on ‘personality neuroscience’, a term denoting the research aimed at neuroimaging correlates of inter-individual temperament and character variability. Among other methods, a graph theoretical analysis of the functional connectivity in resting-state functional magnetic resonance imaging data was applied in a study by Gao et al. (2013), reporting novel functional connectivity correlates of personality traits. The current paper presents a conceptual replication of the results of this study and discusses the related challenges, including an extension of the original statistical methods in order to illustrate the effect of the multiple comparison problem. Five personality dimensions were obtained using the revised ‘Big Five’ Personality Inventory, including scores of Extraversion and Neuroticism covered in the original paper. Using a larger sample (84 subjects) with adequate statistical power (ranging from 0.75 to 0.95 across analyses), we failed to replicate any of the nine specific neuroimaging correlates of personality presented by Gao et al. While acknowledging differences in the experimental procedures, we discuss that the lack of replication might be caused by the relatively liberal control of false positives in the original study. Indeed, the original testing scheme leads to an expected count of about 10 false positive observations among all tests; applying this scheme to our data we observed a similar number of positive tests, albeit for different relations. No significant correlations were found in our data when standard family-wise error control was applied. These results illustrate the importance of combining exploration with independent validation, use of large datasets, as well as appropriate control of multiple comparison problem in order to prevent false alarms in research into neural substrates of personality differences. Importantly, our findings do not disprove the existence of a link between personality and the brain’s intrinsic functional architecture; but rather suggest that such a link might be even more subtle and elusive than previously reported.

## Introduction

For centuries scientists have been interested in human personality in an attempt to understand inter-individual differences and possibly label the unique psychological structure of human behavior using heterogeneous approaches. The dimensional approach is one of the most prominent methods used to describe both normal and disturbed behavioral patterns or personality dimensions, mostly based on standard psychometric questionnaires.

Probably the most popular Big Five Factor model of personality traits is based on factor analysis of verbal descriptors (responses on personality questionnaire) divided into five factors defined as openness to experience, conscientiousness, extraversion, agreeableness, and neuroticism (Costa et al., 1992). Similarly, Eysenck (1991) proposed an alternative three-factor model (psychoticism, extraversion, neuroticism). In fact, the two models have been shown to be strongly related (Scholte and Bruyn, 2004; Zuckerman et al., 1993; Aluja et al., 2002).

Currently, it is broadly accepted that stable interindividual differences in how people think, feel and behave are in some way reflected in the complex brain architecture and dynamics. However, the precise neurobiological substrate of individual personality traits remains unclear. Recently, the number of studies focused on ‘Personality neuroscience’ has been growing. Most of these studies apply neuroimaging in order to find specific personality correlates in the brain anatomy or functionality using a wide range of data analysis methods (Adelstein et al., 2011; Wei et al., 2011; Haas et al., 2007; Wright et al., 2006; Aghajani et al., 2014; DeYoung and Gray, 2009; Lei et al., 2013, 2015; Dubois et al., 2018). Even though these brain localization methods offer a useful tool for testing of some research ideas, a cautious methodological approach has been suggested when similar whole-brain analysis is applied in search for association with any of the measured personality dimensions (Cacioppo et al., 2003; Willingham and Dunn, 2003) in order to prevent false positive findings (Tomeček et al., 2020). These concerns are further underscored by evidence of poor test-retest reliability of common fMRI measures (Elliott et al., 2020). The challenges of reproducibility in neuroimaging have been highlighted by large-scale replication efforts (Open Science Collaboration, 2015) and investigations into inflated false-positive rates in fMRI inferences (Eklund et al., 2016).

In the neuroimaging community, there has been a recent growth of interest in resting-state functional magnetic resonance imaging (fMRI). Despite its simple administration, this method allows the analysis of multiple brain systems from a single acquisition. Thus, the resting-state fMRI easily found its way into personality neuroscience (e.g. Lei et al. (2013); Gao et al. (2013); Lei et al. (2015)). For instance, it was suggested that personality traits (such as extraversion) might be reflected by some aspects of the scaling property and long-range temporal dependence of resting-state networks such as the default-mode network (Lei et al., 2013, 2015).

A key method applied in resting-state fMRI is functional connectivity analysis defined as the temporal correlations between spatially remote neurophysiological events (Friston, 1994). Recently, graph-theoretical approaches have been proposed for analysis of brain network organization and connectivity (Bullmore and Sporns, 2009). Importantly, graph-theoretical measures derived from correlation networks, including small-world properties, may be susceptible to methodological artifacts inherent to the correlation structure itself (Hlinka et al., 2017), an issue that has been systematically explored using network correspondence frameworks (Kong et al., 2025) and through examination of how spatial and temporal autocorrelation affects network inference (Shinn et al., 2023). The graph theory of functional connectivity was used by Gao et al. (2013), who aimed to test the theoretically derived biological personality model proposed by Eysenck. On a sample of 71 subjects they reported that the normalized clustering coefficient values of the whole-brain functional networks are positively correlated with extraversion scores (*r* = 0.345, *p* = 0.004). In addition, they found multiple positive associations with neuroticism on a local network level, interpreting them as supporting Eysenck’s biological personality model. However, based on the findings of both positive and negative correlations of extraversion scores in several brain regions, the authors suggest that the relationship between extraversion and regional arousal is not as simple as that proposed by Eysenck (Gao et al., 2013). Furthermore, they argue that the right lateralization of regions reported with regard to neuroticism provided neurofunctional evidence for the preferential involvement of the right hemisphere in emotions and motivational states.

We note that the study by Gao et al. (2013) was among the earlier applications of graph-theoretical functional connectivity analysis to personality neuroscience, and at the time our replication effort was initiated it represented a natural and accessible target: its methodology was clearly described, and its sample size and statistical approach were amenable to independent validation. Indeed, the data collection and analysis reported here were completed several years ago, predating the broader community discussion on replication challenges in neuroimaging (Marek et al., 2022). We wish to emphasize that the statistical concern raised here is not unique to Gao et al. (2013): the multiple comparison problem is pervasive in the personality neuroscience literature, and similar issues are likely present in other studies as well. However, systematically verifying this across the broader literature — particularly for more recent studies employing complex multi-stage analytical pipelines — falls outside the scope of the present work. To test the specific results reported in the original study, we first ensured that our dataset was sufficiently powered to replicate them. To this end, we used restingstate fMRI data acquired in a sample of 84 healthy subjects (compared to the original sample size of 71 subjects) using a similar scanner setting and a personality questionnaire (the NEO Five-Factor Inventory (NEO-FFI) (Costa et al., 1992)). Notably, this dataset is identical to that used in a previous study by Tomeček et al. (2020), which examined seed-based connectivity between default-mode network seeds and the rest of the brain; the present work employs a complementary graph-theoretical analysis approach to examine topological properties of the full 90-region connectome. This inventory captures, among other dimensions, the Neuroticism and Extraversion scales that are highly correlated to the corresponding scales of the Eysenck Personality Questionnaire.

Regarding the statistical methodology, the procedure for correction for multiple testing in the report of Gao et al. (2013) consists of using the threshold *p* = 1*/*90 instead of the common *p* = 0.05. While this indeed provides some correction, it is likely insufficient when carrying out hundreds of tests. Therefore, to shed light on the validity of the original results, we provide both basic theoretical analysis of the effects of such lenient correction on the expected number of false positive results, as well as demonstrate the effect in a loose replication of the analysis procedure on this similar dataset (with both the lenient and a standard multiple testing correction).

## Materials and methods

### Participants

We collected one session of MRI data from 84 participants (48 men, mean age *±* SD: 30.83 *±* 8.48, three left-handed). The study was approved by the Ethics Committees of NIMH (National Institute of Mental Health in Klecany, Czech Republic) and IKEM (Institute for Clinical and Experimental Medicine in Prague, Czech Republic). All subjects gave written informed consent in accordance with the latest version of the Declaration of Helsinki.

### Personality questionnaires

The NEO Five-Factor Inventory (NEO-FFI) (Costa et al., 1992) was used to assess personality dimensions of neuroticism, extraversion, openness, agreeableness and conscientiousness for each subject. Raw scores were transformed into T-scores (in agreement with Gao et al. (2013)) as

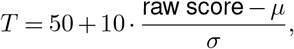

where *μ* represents the mean value of personality scores over all subjects and *σ* is the standard deviation of the scores. The neuroticism and extraversion personality scores obtained from the NEO-FFI questionnaire have been shown to correlate substantially with the corresponding scales of the revised Eysenck Personality Questionnaire used in Gao et al. (2013) (Scholte and Bruyn, 2004; Zuckerman et al., 1993; Aluja et al., 2002), and are considered to capture largely overlapping variance in these personality dimensions. It should be noted, however, that the two instruments differ in their theoretical grounding and item content: the EPQ-RSC neuroticism scale emphasises emotional reactivity and arousal, whereas the NEO-FFI conceptualises neuroticism more broadly, incorporating cognitive and interpersonal facets of negative emotionality. The NEO-FFI also provides finer measurement resolution by virtue of its larger item pool. These differences mean the two scales are related but not fully equivalent psychometric constructs. Crucially, no direct cross-study comparison of personality scores is made in the present work; each study’s neural–personality correlations are computed independently within its own dataset, so score calibration between instruments is not required.

### Image acquisition

All brain scans were obtained using Siemens TrioTim 3T MR machine located at IKEM. Firstly, a high-resolution 3D anatomical T1-weighted image was acquired (TR = 2300ms, TE = 4.63ms, flip angle = 10^*?*^, FOV 256 x 256, image matrix size 256 x 256, voxel size = 1 x 1 x 1 mm, 224 sagittal slices). Then, the functional T2-weighted images with blood oxygenation level-dependent (BOLD) contrast, (TR = 2500 ms, TE = 30 ms, flip angle = 90^*?*^, FOV 192 x 192, image matrix size 64 x 64, voxel size = 3 x 3 x 3 mm, 44 axial slices, 240 volumes in total for each subject) were collected using the echo-planar imaging (EPI) technique. Participants were instructed to lie still with their eyes closed during the resting-state scan.

### Data preprocessing

We used the CONN toolbox (CONN 15.e) in Matlab environment (Matlab 2015b) to perform the preprocessing of functional images, and to calculate the functional connectivity for each subject. The CONN toolbox uses standard SPM (SPM8) modules. As the first step in our preprocessing pipeline of functional images, a slicetiming correction was performed in order to correct different acquisition times of axial slices, followed by spatial smoothing with a 6 mm FWHM kernel. This was followed by a realignment of all functional volumes to the first volume to minimize a possible head motion generated while acquiring images in the scanner. We assessed the subjects’ head motion by frame-wise displacement (FD, a measure of head motion calculated as the sum of absolute values of six motion parameters: three translational and three rotational displacements) (Power et al., 2012); the *mean ± SD* of FD over subjects was 0.0464 *±* 0.0221, with the FD sequences in no subject exceeding the high-motion threshold *FD >* 0.5 mm (Power et al., 2014). No subjects were therefore discarded from the analysis. Finally, all functional images were normalized into stereotaxic space with resolution 3 x 3 x 3 mm using the MNI 152 standard space template.

The CONN toolbox has implemented CompCor method which extracts time series of white matter and cerebrospinal fluid (this method uses subject-specific ROIs created from segmented anatomical scans) from each subject’s functional images and uses them while denoising the final time series. Average time series of white matter, cerebrospinal fluid and six motion parameters were used in a linear regression to remove their possible confounding effect on average time series from regions of interest. Importantly, global signal regression (GSR) was not applied, as the CompCor approach was used instead to control for physiological noise. The automated anatomical labeling (AAL) atlas (Tzourio-Mazoyer et al., 2002) was used to define 90 regions of interest (ROIs), 45 per hemisphere, from which 90 average time series were extracted in order to calculate functional connectivity between them. The resulting time series were linearly detrended and bandpass filtered using the Butterworth filter (band-pass 0.01 – 0.08 Hz) to remove low-frequency drifts and high-frequency physiological noise. However, it should be noted that physiological confounds such as cardiac and respiratory variations can produce structured spatial patterns resembling neuronal connectivity (Chen et al., 2020).

### Functional connectivity graph

Connectivity matrices represent the Fisher z-transformed Pearson correlation coefficients for all ROI pairs defined by the AAL atlas, yielding a 90 *×* 90 correlation matrix for each subject (Liu et al., 2008).

Formally, the undirected edge *e*_*ij*_ between ROIs *i* and *j* is defined as:

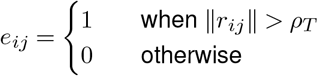

Practically, an edge is assumed between two nodes, if the absolute value of *r*_*ij*_ exceeds a predefined threshold *ρ*_*T*_.

### Graph-theoretical analysis

The following nodal topological network properties were calculated (reflecting Gao et al. (2013)): the degree *K*_*i*_, the clustering coefficient *C*_*i*_, the minimum path length *L*_*i*_, the efficiency *E*_*i*_ and the betweenness centrality *BC*_*i*_ for each node *i*. The global network measures included the average degree *K*, the local network efficiency *E*_*local*_, the global network efficiency *E*_*global*_, the characteristic path length *L*, the clustering coefficient *C*, the normalized versions of the latter two (*λ* and *γ* respectively), and the small-worldness *σ*.

#### Degree

The degree of each node, *K*_*i*_, is simply the number of direct neighbors of a node *i* (Bullmore and Sporns, 2009), implying that the node with a higher degree has more connections. The average degree *K* is the mean of *K*_*i*_ of all nodes.

#### Clustering coefficient

The clustering coefficient *C*_*i*_ quantifies how close the node’s neighbors are to being a clique, representing the ratio of existing connections to all possible connections in the graph. Formally,

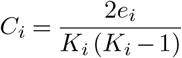

where *e*_*i*_ is the number of edges in the graph (Bullmore and Sporns, 2009; van den Heuvel et al., 2009). The clustering coefficient *C* is the mean of *C*_*i*_ of all nodes.

#### Minimum path length

The minimum path length per node, *L*_*i*_, is defined as the mean shortest absolute path length of node *i* to all other nodes in the graph (Bullmore and Sporns, 2009; van den Heuvel et al., 2009). This quantifies the level of routing efficiency, thus the capability of information propagation in the network. It is defined as

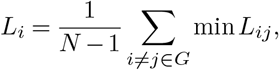

where *L*_*ij*_ is the shortest absolute path length between nodes *i* and *j*, and the absolute path length is the number of edges included in the path connecting the two nodes. The characteristic path length *L* is the average of *L*_*i*_ for each node *i* in the graph.

#### Efficiency

The nodal efficiency *E*_*i*_ is the inverse harmonic mean of the length between node *i* and all other nodes in the graph (Latora and Marchiori, 2001; Bassett and Bullmore, 2006), therefore

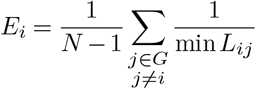

The global efficiency, *E*_*global*_, is the mean of *E*_*i*_ of all the nodes in the network. In the subgraph *G*_*i*_, one can calculate the local efficiency of node *i* as

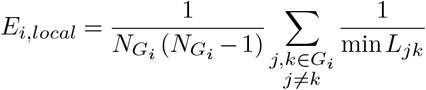

The local efficiency of the network is then similarly the mean of *E*_*i,local*_ of all the nodes in the network (Rubinov and Sporns, 2010).

#### Betweenness centrality

The betweenness centrality of a node, *BC*_*i*_, is defined as a fraction of all shortest paths in the network that pass through node *i* (Rubinov and Sporns, 2010). This centrality measure describes the central nodes that participate in many shortest paths within a network and act as important hubs for the information flow (Freeman, 1978). Mathematically,

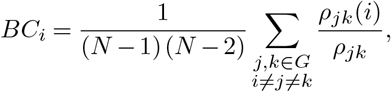

where *ρ*_*jk*_ is the number of shortest paths between nodes *j* and *k*; *ρ*_*jk*_(*i*) is the number of shortest paths between *j* and *k* that pass through node *i*.

#### Small-world parameters

One of the most studied properties of the brain networks is their small-worldness. While random networks exhibit a low clustering coefficient and a typical short path length, the highly organized network (such as full lattice graph) exhibit the opposite — high clustering coefficient, but on average also long path lengths. The network with a small-world organization would show higher clustering than the random graph and similar path length, i.e. *γ* = *C/C*_*random*_ *>* 1 and *λ* = *L/L*_*random*_ ≈ 1 (Watts and Strogatz, 1998). These conditions can be evaluated by the small-worldness coefficient, *σ* = *γ/λ*, and the network is said to be small-world if *σ* ≫ 1 (Humphries et al., 2006). The clustering coefficient and characteristic path length of random graph, *C*_*random*_ and *L*_*random*_, were computed as the average from a set of 100 realizations of random networks with the same degree distribution as that of the examined network. The random networks were generated using the Maslov–Sneppen rewiring algorithm (Maslov and Sneppen, 2002).

Recalling eq. Eq. (), the threshold *T* is defined as a ratio of total number of edges in a graph to maximum possible number of edges (Achard and Bullmore, 2007). In this study, we investigated topological properties of brain network over a range of *T*_*min*_ ≤ *T* ≤ *T*_*max*_ defined in the next section.

### Association between network organization and personality dimensions

In this study, we estimated both global and nodal graph-theoretical measures. All the nodal and global measures were computed for repeatedly thresholded graphs over the range of 0.1 ≤ *T* ≤ 0.31 with an interval of 0.01 (in accordance with Gao et al. (2013)). Then, the area under curve (AUC, summed values of the graph measure across the threshold range) for each network metric was calculated, providing scalar metric for topological characterization of brain network, independent of single threshold selection. As for the association with personality dimensions, the partial correlation was calculated between the AUC of each network metric and personality dimension scores, with age and gender being covariates.

### Statistical inference

As we mentioned in the introduction, the original study used a relatively lenient statistical threshold. We replicated the analysis using both the original thresholding and a more appropriate standard correction procedure as explained below. In order to assess statistical significance of the results we utilized in both cases a permutation test with 20000 permutations for the global network properties and 42000 permutations for the nodal network properties (the number of permutations differs due to minimal number of permutations required for the multiple testing correction). The partial correlation between global and nodal AUCs were recalculated for each permutation of the data, yielding the null hypothesis distribution.

For the ‘replication’ analysis, the percentile threshold for significance was selected as the inverse of number of regions, thus 1*/*90. Note that this threshold, that is without further multiple testing correction prone to provide on average 1 false positive for every 90 tests, was used in accordance with the original study by Gao et al. (2013), who adopted it from Lynall et al. (2010), where it was used in a different context.

As a second, appropriately conservative statistical approach, we have also applied a standard family-wise error (FWE, controlling the probability of any false positives in the set of hypotheses at 0.05) rate controlling Holm’s step-down procedure (Holm, 1979). This method guarantees control of the probability of any false positives in the set of hypothesis at 0.05 by a closed testing procedure. As a side note: similar and potentially better known is the Benjamini–Hochberg step-up procedure (Benjamini and Hochberg, 1995) that controls the false discovery rate (FDR) at 0.05 under some (albeit mild) assumptions. It gave in our case equivalent results.

Finally, we performed a power analysis to assess whether our sample size was sufficient to detect the effect sizes reported by Gao et al. (2013), had those effects been genuine, *at the same per-test significance threshold used in the original study* (*α* = 1*/*90). According to Algina and Olejnik (2003), the distribution of a partial correlation coefficient is the distribution of a correlation coefficient with a sample size reduced by the number of control variables. Therefore, in order to calculate the needed sample size we may use the classic formula for estimating the correlation sample size (e.g. Hulley et al. (2013))

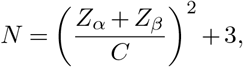

where *Z*_*α*_ and *Z*_*β*_ are the values of the standard normal variable corresponding to *α* (type I error rate) and *β* (type II error rate), respectively, and *C* = 0.5 · ln[(1 + *r*)*/*(1 − *r*)] is Fisher’s arctanh transformation with *r* being the sample correlation coefficient.

Assuming the effect sizes observed in the original study — in particular partial correlation 0.345 for the global graph property (normalised clustering coefficient) against extraversion and 0.318–0.412 for the nodal graph properties — and evaluating power at the original study’s per-test threshold *α* = 1*/*90 ≈ 0.011 (one-tailed), our sample of *N* = 84 provides power 0.83 for the global property and powers 0.75–0.95 for the nodal properties. This confirms that, had the effects reported by Gao et al. (2013) been real, we would have had adequate power to detect them using their own significance criterion.

## Results

### Statistics of the personality dimensions

Table 1 describes the raw scores of the five personality dimensions from NEO Five-Factor Inventory questionnaire, while the Table 2 shows correlations across the scores of the five personality dimensions. Extraversion and neuroticism, the two basic personality measures concerned in this study, show moderate negative correlation (*r* = *−*0.368 with *p* = 0.0005). In agreement with the original study, we added extraversion (resp. neuroticism) scores as covariate (along with age and gender) when calculating the partial correlation between neuroticism (resp. extraversion) and the AUC of each network metric. In the subsequent analysis of the remaining three personality variables unavailable to the original study, we added other personality measures which showed significant correlation (*p <* 0.05) as covariates.

**Table 1.** Descriptive statistics of the five personality dimensions of 84 participants. Age and z-scored personality scores shown as mean *±* SD.

| Category | Data |
| --- | --- |
| gender (male / female) | 48 / 36 |
| age (years) | 30.83 ± 8.48 |
| neuroticism (N) | -0.369 ± 1.117 |
| extraversion (E) | 0.085 ± 0.858 |
| openness (O) | 0.543 ± 1.075 |
| agreeableness (A) | 0.284 ± 0.876 |
| conscientiousness (C) | 0.274 ± 0.888 |

**Table 2.** Correlations between scores of the five personality dimensions: **N** — neuroticism, **E** — extraversion, **O** — openness, **A** — agreeableness and **C** — conscientiousness. Asterisk denotes *p <* 0.05.

|  | N | E | O | A | C |
| --- | --- | --- | --- | --- | --- |
| N | X | -0.368 ($p = 0.0005^*$) | 0.174 ($p = 0.114$) | -0.272 ($p = 0.012^*$) | -0.399 ($p = 0.0002^*$) |
| E | | X | 0.115 ($p = 0.295$) | 0.372 ($p = 0.0005^*$) | 0.264 ($p = 0.0152^*$) |
| O | | | X | 0.109 ($p = 0.325$) | -0.203 ($p = 0.065$) |
| A | | | | X | 0.184 ($p = 0.0933$) |

### Validation of the original study

The first step in this study was to compare the results of Gao et al. (2013) with our findings. Table 3 summarizes results in terms of partial correlation and p-value of both studies.

**Table 3.** Significant results of partial correlation between global and nodal graph-theoretical measures and personality dimension scores from Gao et al. (2013) study compared to results from our study using the same statistical threshold.

| dimension | AUC of | ROI | Gao et al. (2013) |  | this study |  |
| --- | --- | --- | --- | --- | --- | --- |
|  |  |  | partial corr. | p-val | partial corr. | p-val |
| extraversion | $\gamma$ | global measure | 0.345 | 0.004* | -0.115 | 0.308 |
| extraversion | $BC_i$ | insula L | 0.318 | 0.008* | 0.131 | 0.235 |
| extraversion | $BC_i$ | middle temporal gyrus L | -0.412 | 0.0005* | 0.030 | 0.783 |
| extraversion | $BC_i$ | middle temporal gyrus R | -0.337 | 0.005* | 0.176 | 0.114 |
| neuroticism | $BC_i$ | precentral gyrus R | 0.363 | 0.002* | -0.122 | 0.292 |
| neuroticism | $BC_i$ | olfactory cortex R | 0.319 | 0.008* | 0.111 | 0.350 |
| neuroticism | $BC_i$ | amygdala L | 0.384 | 0.001* | -0.120 | 0.293 |
| neuroticism | $BC_i$ | amygdala R | 0.370 | 0.002* | -0.067 | 0.553 |
| neuroticism | $BC_i$ | caudate nucleus R | 0.314 | 0.009* | -0.059 | 0.619 |

As clearly seen from the results in Table 3, we did not reproduce any of the associations between graph-theoretical measures and neuroticism or extraversion reported in the original study.

### The associations between global network metrics and personality dimensions

In this part of the study we carry out exploratory testing of relations of all the considered global and nodal network properties with neuroticism and extraversion personality dimensions. To illustrate the effect of multiple comparisons treatment, we carry out the analysis using two different statistical procedures. The first is using the fixed significance threshold *p* = 1*/*90 as done by Gao et al. (2013); the second is controlling the family-wise error rate by the Holm’s step-down procedure (Holm, 1979).

First, we computed partial correlations between AUC of global network properties and neuroticism and extraversion personality dimensions. When computing the partial correlation, age, gender, and the other dimension were used as covariates. The permutation test was evaluated as follows: we created 20000 random permutations and permuted the AUC of global metrics, keeping all the personality scores and covariates unpermuted. From the ensemble of permutations we evaluated the null distribution for each AUC of network metric and computed the two-tailed p-value. The results are summarized in Table 4. Since no significant correlation at *p <* 0.05 was found, the multiple testing procedure was not even necessary to employ.

**Table 4.** Partial correlations between AUC of global network metrics and two personality measures: neuroticism and extraversion. Two-tailed p-values computed using permutation test with 20000 random permutations.

| *AUC of* | neuroticism |  | extraversion |  |
| --- | --- | --- | --- | --- |
|  | *partial corr.* | *p-val* | *partial corr.* | *p-val* |
| char. path length | 0.1236 | 0.2690 | 0.0713 | 0.5095 |
| local efficiency | -0.0400 | 0.7020 | -0.1440 | 0.2005 |
| global efficiency | -0.0842 | 0.4670 | -0.0900 | 0.4125 |
| avg. clustering | 0.1236 | 0.2710 | -0.0274 | 0.8005 |
| norm. clustering | -0.0310 | 0.8110 | -0.1149 | 0.3075 |
| norm. path length | 0.1373 | 0.2135 | 0.0601 | 0.5705 |
| small-worldness | -0.0655 | 0.5405 | -0.1228 | 0.2745 |

### The associations between nodal network metrics and personality dimensions

Next, the relationship between AUC of nodal network metrics and personality dimensions was sought. As in the former case of global network metrics, firstly we computed the partial correlation for neuroticism and extraversion scores.

When considering neuroticism and extraversion and their relationship with nodal network metrics, we found a small number of regions where the partial correlation is significant at the original level *p* = 1*/*90 used by Gao et al. (2013), as shown in Table 5.

**Table 5.** Significant results of partial correlation between nodal graph-theoretical measures and personality dimension scores before the FWE correction addressing multiple comparison problem.

| *dimension* | *AUC of* | *ROI* | *partial corr.* | *p-val* |
| --- | --- | --- | --- | --- |
| extraversion | $BC_i$ | gyrus rectus L | 0.293 | 0.007* |
| extraversion | $K_i$ | superior frontal gyrus L | -0.326 | 0.002* |
| extraversion | $C_i$ | orbital superior frontal gyrus R | -0.359 | 0.002* |
| extraversion | $C_i$ | angular gyrus R | -0.289 | 0.009* |
| extraversion | $E_i$ | parahippocampal gyrus L | -0.277 | 0.008* |
| extraversion | $E_i$ | angular gyrus R | -0.277 | 0.009* |
| neuroticism | $C_i$ | middle frontal gyrus R | 0.275 | 0.015* |

However, since we are considering a set of statistical inferences simultaneously we are facing a problem of multiple comparisons. In fact, we are inferring 180 statistical significances per nodal metric at the same time, that is 900 correlation in total. To tackle the multiple comparisons problem, we adopted a multiple testing correction procedure. The result of multiple testing correction was that none of the observed correlations are to be considered statistically significant. Therefore we conclude that our study did not uncover any relationship between AUC of network metrics and neuroticism and extraversion scores. In the next section, we explore any potential relationships with the remaining three personality dimensions measured by NEO-FFI.

### The associations between network metrics and openness, agreeableness and conscientiousness

We repeated the analysis and computed partial correlations and their respective p-values between AUC of global network measures and openness, agreeableness (extraversion as covariate) and finally, conscientiousness (with neuroticism as covariate). In all computations, the age and gender were used as covariates as well. The results are summarized in Table 6. As in the former case, no significant correlations were found (even without applying the multiple testing correction procedure).

**Table 6.** Partial correlations between the AUC of global network metrics and three additional personality measures: openness, agreeableness and conscientiousness. The values for neuroticism and extraversion are shown in table 4. Two-tailed p-values computed using permutation test with 20000 random permutations. Since no significant correlation was found, the multiple testing correction procedure was not necessary to employ.

| *AUC of* | openness |  | agreeableness |  | conscientiousness |  |
| --- | --- | --- | --- | --- | --- | --- |
|  | *partial corr.* | *p-val* | *partial corr.* | *p-val* | *partial corr.* | *p-val* |
| char. path length | -0.0811 | 0.4535 | 0.0264 | 0.8385 | -0.1332 | 0.2370 |
| local efficiency | 0.0223 | 0.8425 | 0.1201 | 0.2910 | -0.0652 | 0.5615 |
| global efficiency | 0.2063 | 0.0600 | 0.0852 | 0.4465 | 0.0299 | 0.8010 |
| avg clustering | -0.0962 | 0.4020 | 0.1164 | 0.2935 | -0.1368 | 0.2440 |
| norm. clustering | 0.1030 | 0.3645 | 0.0394 | 0.7255 | 0.0711 | 0.5070 |
| norm. path length | -0.0432 | 0.7025 | 0.0419 | 0.6910 | -0.1418 | 0.2015 |
| small-worldness | 0.0986 | 0.3650 | 0.0270 | 0.8135 | 0.0902 | 0.4405 |

In the case of nodal properties, we found in total 15 significant correlations between AUC of nodal network metrics and personality measures (1 for agreeableness, 3 for conscientiousness, 11 for openness, overall corresponding exactly to 1/90 false positives out of 1350 tests). As before, we employed the procedure to control the FWE rate at 0.05 to deal with the multiple comparison problem and again, none of the correlations were then significant with this correction, suggesting that they occurred randomly as false positives. Again, we conclude that no relationship was observed between the AUC of nodal network metrics and the three additional personality domains.

## Discussion

In the presented study, we did not replicate the results of the original study by Gao et al. (2013). Moreover, our results suggest that the observed significant partial correlations between personality dimensions and network metrics are most likely accidental. Our evidence for this is threefold. Firstly, we have shown that the original testing procedure is prone to generate multiple false positive results. Specifically, the report by Gao et al. (2013) uses a fixed significance threshold *p* = 1*/*90 throughout — a threshold that is prone to providing on average 1 false positive in 90 tests. Therefore, particularly when testing the nodal properties, out of the 5 × 2 × 90 = 900 tests carried out, on average 10 false positives are to be expected, making it conceivable that a substantial proportion of the 9 reported significant relations constituted false positives; for broader context, we refer the reader to the discussion by Ioannidis (2005) on why most published research findings are false. Secondly, our study, conducted on a sample of comparable size to the original, did not replicate any of the previously reported 9 FC-graph correlates of personality - instead, with the same procedure we observed a completely disjoint set of other (likely false positive) results, that did not survive an appropriate multiple testing procedure. Finally, Holm’s step-down procedure for dealing with the multiple testing problem deals with the smallest p-values in increasing order (note that Holm’s method is a step-down procedure, distinct from the Benjamini–Hochberg step-up procedure for FDR control), we can reproduce what would have been the results of its application on the original data. In particular, the minimum p-value observed in the original study was 0.0005, while the adjusted FWE threshold for 900 tests was one order smaller.

When applying an appropriate multiple testing procedure it becomes clear, that the amount of hypotheses about the *nodal* properties becomes a critical factor decreasing the study power. Based on this we would recommend focusing future studies on global network properties, or hypothesis-driven subselection of nodal properties of specific brain areas, to limit the number of tests taken. In general, limiting the number of hypotheses is key for keeping reasonable power even for larger sample sizes, although alternative dimension reduction techniques or statistical approaches might also help with this situation in network theory applications; this is a matter of ongoing methodological research. Alternatively, the tests of nodal properties can be considered exploratory and requiring independent validation. From this viewpoint, our study did not confirm the hypotheses generated by the previous report by Gao et al. (2013).

Some limitations of our study should be mentioned in order to present a complete picture. Even though the analysis procedure applied in our study was the same as in the original research, we admit that some methodological differences could account for discrepancies in the obtained results (albeit many of the originally reported results are likely false positives due to purely statistical reasons). Compared to Gao et al. (2013), the following methodological alterations should be mentioned in particular: NEO-FFI administration instead of the Eysenck Personality Questionnaire and slightly different scanning sequence parameters. Therefore, the experimental part of this work does not represent a replication study in the strictest sense, but rather a conceptual validation using slightly different tools. The substitution of the EPQ-RSC with the NEO-FFI was motivated by its availability in our data collection protocol, which was designed as part of a broader neuroimaging project. Both models share strongly correlated factors of extraversion and neuroticism (Scholte and Bruyn, 2004); moreover, Eysenck’s factor of psychoticism has been shown to relate (negatively) to agreeableness and conscientiousness (Zuckerman et al., 1993). The moderate negative correlation between extraversion and neuroticism observed in our sample (*r* = − 0.368, *p* = 0.0005) is in agreement with prior studies (Rusting and Larsen, 1997; Wright et al., 2006), supporting the representativeness of our cohort. Nevertheless, as discussed in the Methods, the two instruments are related but not fully equivalent psychometric constructs: they differ in theoretical grounding, item content, and measurement sensitivity. It therefore cannot be excluded that a portion of the observed differences in results is attributable to the instrument substitution rather than — or in addition to — the statistical issues in the original study. We consider this an inherent limitation of any conceptual replication, and one that is difficult to resolve without a dataset containing both instruments administered to the same subjects. Further, concerning the MRI data, we used the same scanner as the original study, however with different sequence parameters. Nevertheless, the settings were relatively similar – the main differences were in using a slightly longer TR (2.5s instead of 2s), overall acquisition time (600s vs 510s), and smaller voxel size (3 mm isotropic instead of 3.75 × 3.75 × 4 mm). In general, we argue that these differences in assessment methods are not likely responsible for the observed differences in results, although such methodological variations can generally affect study sensitivity.

It is important to note that, based on the observed results and methodological analysis of the original study, we do not and cannot take the position that the association between brain architecture and personality dimensions does not exist. Rather, based on the results presented above, we only recommend caution in statistical processing of the whole brain with multiple comparisons. This caution is strongly supported by recent large-scale evaluations of brain-wide association studies (BWAS), such as Marek et al. (2022), which demonstrated that reproducible associations between inter-individual differences in brain function and complex phenotypes typically require sample sizes in the thousands—far exceeding those available in standard neuroimaging studies. The poor test-retest reliability of fMRI measures further underscores these replication challenges (Elliott et al., 2020). Also, it could be proposed that some alternative approach to the graph theory of functional connectivity networks might be more beneficial in order to obtain robust and reproducible results. In particular, the interpretation of small-world properties in correlation networks should be treated with caution, as these may reflect inherent properties of correlation matrices rather than genuine underlying network organization (Hlinka et al., 2017).

One possible approach lies in avoiding the multiple testing problem by extracting only one or a small number of strong summary features, such as applied in different contexts like the prediction of personality from fMRI during naturalistic viewing conditions (Jajcay et al., 2025), or prediction of disease symptoms in multiple sclerosis (Rehák Bučková et al., 2022). Another approach is the application of heavily multivariate machine learning approaches to whole brain connectivity patterns. Successful examples of this can be found in largescale studies utilizing the Human Connectome Project (HCP) dataset. For instance, Cai et al. (2020) utilized connectome-based predictive modeling to successfully predict four of the five personality factors, though they noted that controlling for confounds reduced the robustness of these predictions. Similarly, Dubois et al. (2018) reported that rs-fMRI measurements carry information about intelligence and the Big Five factor of openness to experience. However, these recent studies also demonstrated that no single anatomical structure or network was responsible or necessary for the prediction; instead, this prediction relied on redundant information distributed across the brain. We refer the reader to these papers for a detailed discussion of the challenges in connectome-based predictive modelling in personality neuroscience.

## Conclusions

The main goal of this study was to provide additional evidence concerning the reported relationships between personality and the brain’s dynamical repertoire as quantified by graph theory applied to functional connectivity analysis of resting-state and presented by Gao et al. (2013). We conclude that we were not able to validate the specific relationships using an independent dataset; conjecturing that (many of) the original results may constitute false positive findings (Tomeček et al., 2020), as the original study contained relatively liberal control for multiple comparisons, and none of the results was reproduced (even at uncorrected p-level) in our independent study of comparable sample size using closely related measurement tools. Our exploratory analysis for the other possible connections between FC properties and personality using an appropriate correction for multiple testing did not find any significant associations, however such exploration would likely require much larger samples for reasonable power. The presented evidence suggests that validation on independent samples, use of large datasets, and/or stringent control of multiple comparison problem should be meticulously applied in order to control false alarms in research into neural substrates of personality differences.

